# Chondroitin sulfates enhances the barrier function of basement membrane assembled by heparan sulfates

**DOI:** 10.1101/2022.01.23.477428

**Authors:** Chenqi Tao, Neoklis Makrides, Yihua Wu, Steven E Brooks, Jeffrey D. Esko, Xin Zhang

## Abstract

Glycosaminoglycans (GAGs) are ubiquitously expressed polysaccharides attached to proteoglycans, but their functions in the retina are poorly understood. Here we generated conditional knockouts of biosynthetic enzymes for heparan sulfate (HS) and chondroitin sulfate (CS) in retinal progenitor cells. We showed that ablation of HS polymerase Ext1 did not affect initial progression of retinal angiogenesis, but it disrupted the pruning of blood vessels and establishment of arterioles and venules. In the absence of retinal HS, blood vessels were also vulnerable to high oxygen tension in early postnatal stages, which can be rescued by exogenous VEGF, consistent with the role of retinal HS in the fine-tuning of VEGF signaling. Furthermore, we observed that the retinal inner limiting membrane (ILM) was disrupted by deletion of *Ext1* in a timing specific manner, suggesting that retinal HS is required for the assembly but not the maintenance of the ILM. Lastly, we showed that further deletion of C4st1, a CS sulfation enzyme, did not affect the assembly of the ILM, but aggravated the ILM permeability when combined with *Ext1* deletion. These results demonstrated an important role of CS and HS in establishing the barrier function of basement membrane.

## Introduction

Glycosaminoglycans (GAGs) are a family of ubiquitously expressed polysaccharides that play important roles in development and physiology (Bishop et al., 2007; Mikami and Kitagawa, 2013; Mizumoto et al., 2013). Based on their composition, these macromolecules are classified into heparan sulfate (HS), chondroitin sulfate (CS), dermatan sulfate (DS) and keratan sulfate (KS). GAG synthesis require cascades of enzymes, including enzymes involved in the generation of nucleotide sugar precursors and enzymes such as Chsy/Chpf and Ext1/2 that assemble the backbone of CS and HS chains, respectively (Fig. 1A). During assembly of the chains, they undergo various sulfation reactions. In CS, these reactions are catalyzed by members of the Chst family of enzymes that add sulfate groups to the C4 and/or C6 positions of N-acetylgalactosamine residues and to iduronic acid residues in DS. In HS, Ndst isozymes catalyze the initial N-deacetylation and N-sulfation of N-acetylglucosamine residues, which is then followed by the addition of sulfate groups to C2, C6 and C3 of glucosamine residues and to C2 of uronic acids. The extent of these modifications varies, creating enormous structural complexity, which in turn tunes the selectivity of protein binding and subsequent biological activity.

**Figure 1.**
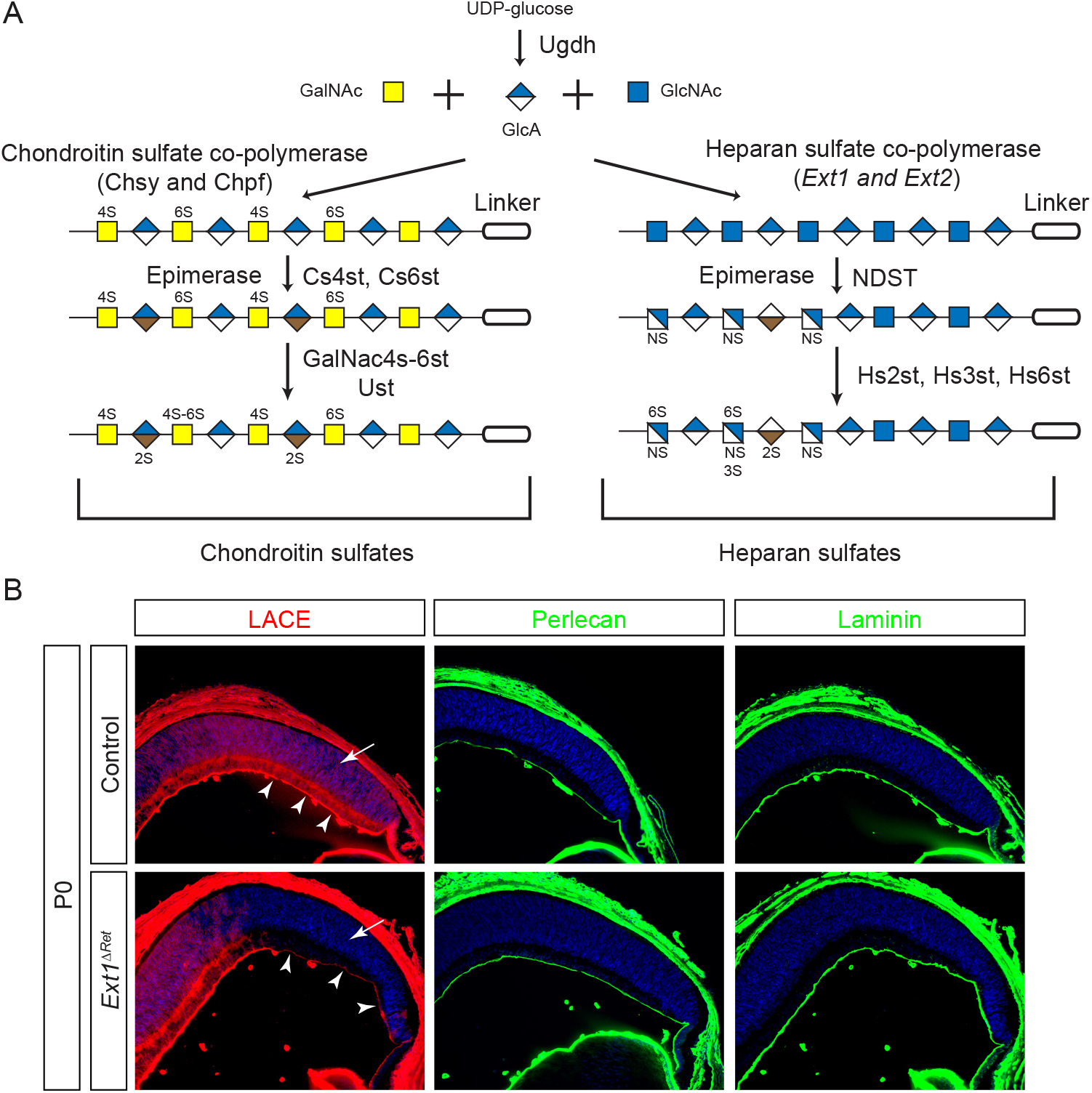
Genetic deletion of *Ext1* abolished HS biosynthesis in the retina. **(A)** The starting monosaccharides for GAG synthesis are Glucuronic acid (GlcUA) produced by Ugdh and N-acetylgalactosamine (GalNAc) for CS or N-Acetylglucosamine (GlcNAc) for HS. They were concatenated by CS co-polymerases Chsy and Chpf or HS co-polymerase Ext1 and Ext2, respectively. Sulfation of CS at the 4-O and 6-O positions were catalyzed first by Cs4st and Cs6st and later by GalNac4s-6st and Ust. In contrast, HS was first sulfated by NDST, followed by epimerization and sulfation at 2-O, 3- and 6-O positions by Hs2st, Hs3st and Hs6st. **(B)** In *Ext1*^*ΔRet*^ mutants, LACE assay showed that HS was efficiently depleted in the peripheral neural retina (arrows), but there was still residual HS in the ILM layer (arrowheads). The ILM also maintained normal patterns of Perlecan and Laminin staining.

Retinal angiogenesis is a prime example that illuminates the diverse functions of GAGs in development. In mice, formation of the retinal vasculature requires sequential migration of astrocytes and endothelial cells into the neonatal retina; both processes are dependent on retinal GAGs (Tao and Zhang, 2014). We have previously examined the impact of altering the enzyme Ugdh, which is responsible for the formation of UDP-glucuronic acid required for the assembly of CS and HS. Inactivation of Ugdh disrupted the inner limiting membrane (ILM) of the retina, which serves as the substratum for astrocyte migration (Tao and Zhang, 2016). HS has also been shown to interact with Vegf produced by astrocytes, generating localized Vegf concentration gradients to attract the migration of endothelial cells into the retina (Ruhrberg et al., 2002; Stalmans et al., 2002). Mutation of the HS interaction motif in Vegfa led to severe defects in vascular outgrowth and patterning. Therefore, migration of retinal astrocytes and endothelial cells requires the distinct functions of GAGs in basement membrane and VEGF signaling.

In this study, we further explored the roles of GAGs in retinal angiogenesis and astrocyte migration by genetically disrupting the biosynthesis of HS and CS. Surprisingly, unlike the profound angiogenesis defect observed in *Vegfa* mutants lacking HS interactions, deletion of HS polymerase *Ext1* in the retina fails to perturb the initial expansion of the vasculature. Instead, loss of retinal HS only impaired vessel remodeling and maturation. On the other hand, the ILM was disrupted by deletion of Ext1 in a timing dependent manner, suggesting that HS is required for the initial assembly but not maturation of the ILM. Lastly, we show that loss of CS sulfation enzyme C4st1 (*Chst11*) did not cause obvious retinal phenotype, but it synergized with *Ext1* deletion in breaching the barrier function of the ILM. These results reveal distinct requirements of HS and CS in the assembly and function of retinal basement membrane.

## Results

### Retinal angiogenesis requires HS for the remodeling but not the outgrowth of the vasculature

Despite the widely accepted role of HS in VEGF signaling, the function of retinal HS in angiogenesis has not been directly tested. To this end, we abolished HS synthesis by ablating the HS polymerase gene *Ext1* using *α-Cre*, which is expressed in the peripheral retina beginning at E10.5 (Cai et al., 2011; Marquardt et al., 2001). We first performed the ligand and carbohydrate engagement (LACE) assay to confirm the loss of HS. In this experiment, the presence of HS on tissue sections was detected by exogenously applied FGF10 and FGFR2b protein, which together bind avidly and specifically to sulfated HS (Pan et al., 2008). As expected, the LACE signal was observed throughout the control eye at postnatal day 0 (P0), most prominently in the ILM overlying the retina (Fig. 1B, arrowheads). In contrast, *α-Cre;Ext1*^*flox/flox*^ (*Ext1*^*ΔRet*^) mutants displayed specific loss of LACE signals in peripheral retinae (Fig. 1B, arrow), which corresponded to the known pattern of *α-Cre* expression. Interestingly, the LACE staining was reduced but not eliminated in the ILM above the mutant area (Fig. 1B, arrowheads), whereas the staining of Perlecan, a major heparan sulfate proteoglycan (HSPG) in the basement membrane, was unaffected. The integrity of the ILM was further confirmed by the continuous staining of Laminin in *Ext1*^*ΔRet*^ mutants, suggesting that genetic ablation of *Ext1* abrogated HS synthesis in the retina without abolishing the ILM formation.

We next examined whether depletion of HS affected retinal angiogenesis, which begins with IB4-positive endothelial cells emerging from the optic disc in the neonatal mouse before spreading to the periphery of the retina (Fig. 2A). Between control and *Ext1*^*ΔRet*^ mutants, we did not observe any statistically significant difference in the outgrowth of retinal vasculature. Nevertheless, the pruning of nascent vessels was significantly altered in *Ext1*^*ΔRet*^ mutants, as shown by the increasing number of empty vessel sheathes positive for the basement membrane marker Col IV, but negative for the endothelial marker IB4 (Fig. 2B, arrows). SMA staining further revealed fewer arterioles and venules in three weeks old *Ext1*^*ΔRet*^ mutant retinae compared to wild-type (Fig. 2C). These results showed that the retinal *Ext1* was dispensable for initial retinal angiogenesis but necessary for the remodeling of the vasculature.

**Figure 2.**
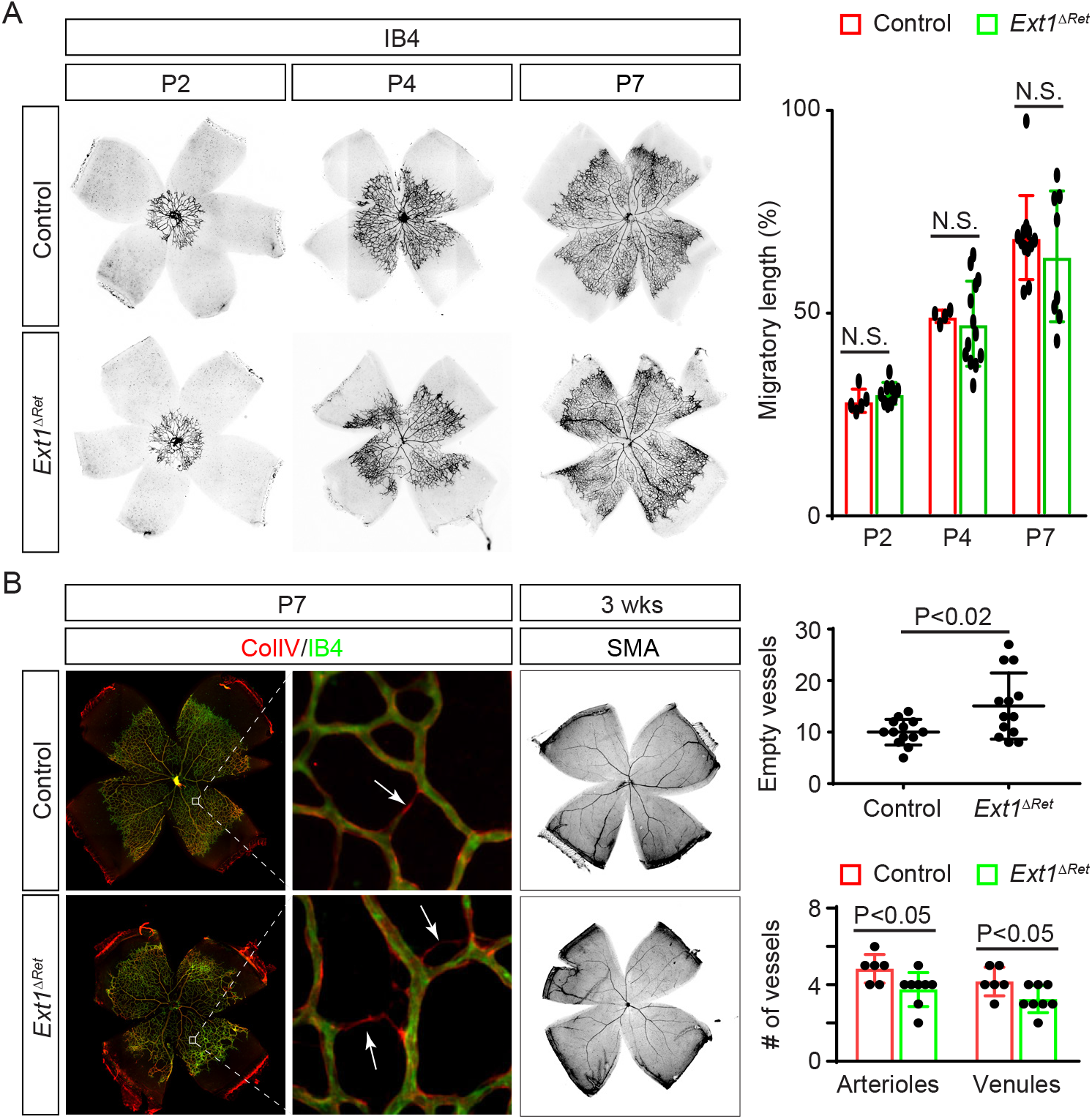
Retinal HS was dispensable for initial angiogenesis but required for vascular pruning. **(A)** The progression of the vascular front was unaffected in *Ext1*^*ΔRet*^ mutant retinae. This was quantified by measuring the migratory distance of endothelial cells in the front of the vascular plexus during postnatal development and normalized as the percentage of the radial length of the retina (Student’ t test, *n*>4 for each sample. N.S., not significant). **(B)** There was a significant increase in the number of Col IV^+^/IB4^−^ empty vessel sheathes in *Ext1*^*ΔRet*^ mutant retinae at P7 (arrows). In contrast, the number of arterioles and venules distinguishable by SMA staining were reduced. Empty vessel sheathes were counted in the middle region of the retina, while arterioles and venules were counted at the optic disc (Student’ t test, *n*=13 for empty sheathe measurement, P<0.02; *n*>6 for vessel measurement, P<0.05).

### Retinal HS is required for VEGF dependent vessel maturation

The relatively mild vasculature defects in *Ext1*^*ΔRet*^ mutant retinae raise questions as to the role of HS in VEGF signaling. In addition to inducing angiogenesis, VEGF signaling also promotes maturation of blood vessels, making them less vulnerable to a hyperoxic environment. To test this function of VEGF signaling in *Ext1*^*ΔRet*^ mutants, we moved P9 pups from the room air and housed them in a 75% O_2_ environment for 7 days (Fig. 3A). In control animals, this treatment led to an expected loss of capillaries in central retinae, leaving major vessels intact (Fig. 3B). In contrast, *Ext1*^*ΔRet*^ mutants exhibited far more severe vessel dropout, extending to include the peripheral retinae, and even included arterioles and venules. If we postponed the hyperoxia treatment until P24, however, both control and *Ext1*^*ΔRet*^ mutant retinae were able to maintain intact vessel plexus, consistent with the eventual maturation of their retinal vasculatures. Therefore, the retinal specific deletion of *Ext1* delayed but did not prevent vessel maturation. We reasoned that if the delayed vessel maturation in *Ext1*^*ΔRet*^ mutants is due to impaired VEGF signaling, it may be ameliorated by exogenous Vegfa. To test this hypothesis, we injected VEGF-A into the vitreous of the eye at P9 prior to the hyperoxia treatment, which unlike VEGF-A injection at P7, did not substantially affect the extent of vessel ablation in wild type animals (Alon et al., 1995). In *Ext1*^*ΔRet*^ mutants, however, intravitreal injection of VEGF-A led to a significant rescue of microvasculature compared to their saline-injected controls (Fig. 3B). These results support that subtle impairment of VEGF signaling plays a role in the vasculature defects in *Ext1*^*ΔRet*^ mutants.

**Figure 3.**
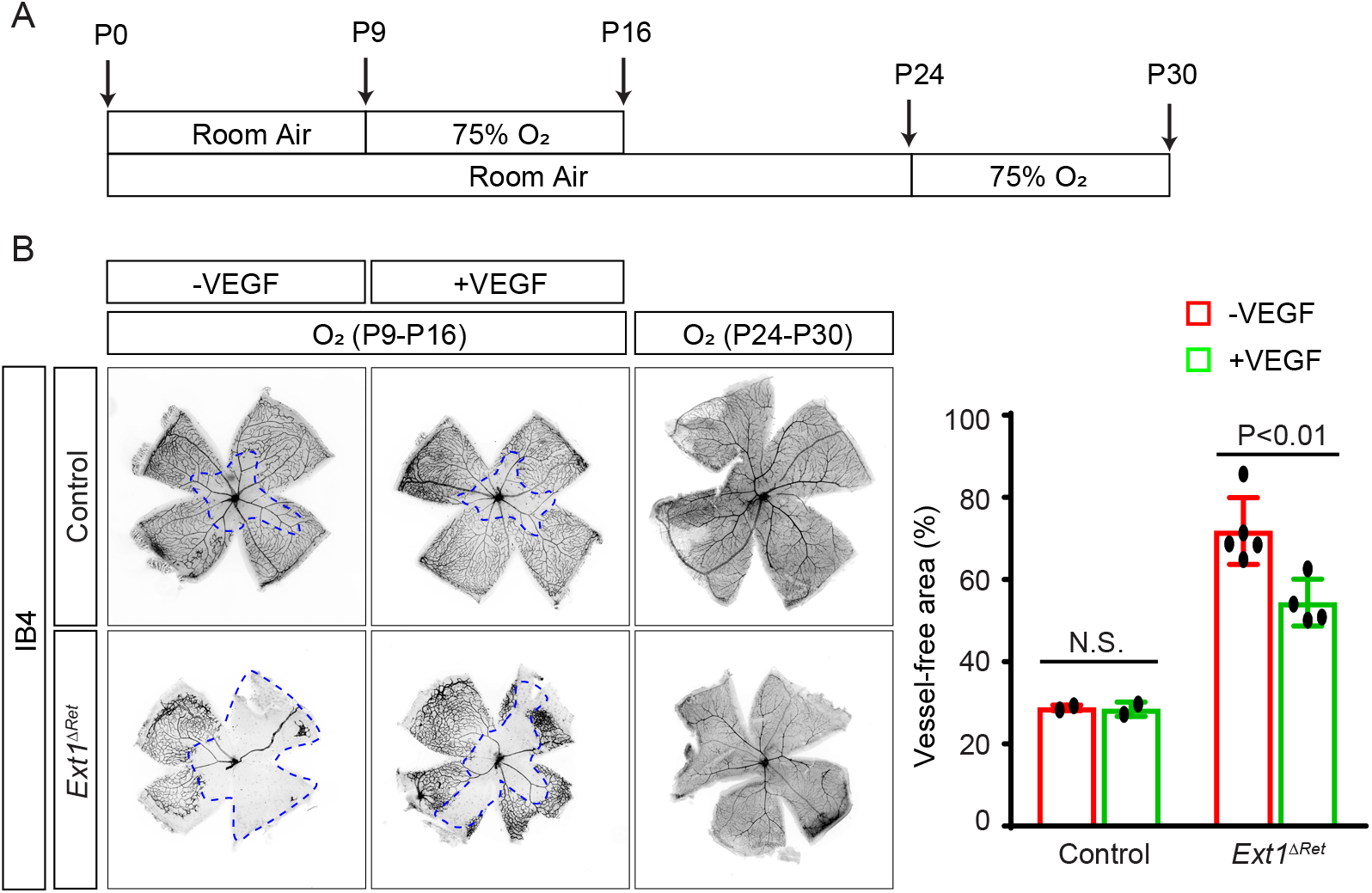
VEGF ameliorated the vessel maturation defect caused by the loss of retinal HS. **(A)** Schematic diagram of the hyperoxia regime. **(B)** Vessel ablations caused by hyperoxia treatment starting at P9 were significantly larger in *Ext1*^*ΔRet*^ mutant retinae that those of controls, which were reduced by intravitreal injection of VEGF-A (70 ng per eye). If the mice were placed in the hyperoxia environment from P24, neither control nor *Ext1*^*ΔRet*^ mutant displayed any obvious loss of vasculature. The area of vessel ablation was measured as percentage of the entire retina (Student’ t test, *n*=2 for each control group, *n*>4 for *Ext1*^*ΔRet*^ control mutant group. N.S., not significant, P<0.01).

### Deletions of HS biosynthesis genes disrupted astrocyte development in the distal retina

We have previously reported severe astrocyte migration defects in *α-Cre;Ugdh*^*flox/flox*^ mutants, whose biosynthesis of HS and CS were abolished (Tao and Zhang, 2016). This prompted us to examine whether similar defects could be observed in HS mutants. At P4, astrocytes had reached the peripheries of control retinae, but the migratory front of astrocytes appeared uneven in *Ext1*^*ΔRet*^ mutants (Fig. 4A, dotted lines). As shown by Pax2 staining in control retinae, nuclei of migrating astrocytes typically displayed spindle shapes, but in *Ext1*^*ΔRet*^ mutants, astrocytes nuclei were more rounded (Fig. 4A, inserts). In contrast, cell bodies of astrocytes labeled by Pdgfra were hyperplastic in *Ext1*^*ΔRet*^ mutants, resulting in a denser astrocytic network. At the end of migration, maturing astrocytes wrap around IB4-positive blood vessels and express the glial cell marker GFAP, but heightened expression of GFAP is also a hallmark of gliosis and Müller cell activation. Compared to the even expression of GFAP in the control retina at P17, *Ext1*^*ΔRet*^ mutants displayed significant increases in GFAP expressions in peripheries of retinae (Fig. 4A), suggesting astrocytes in these regions were under stress.

**Figure 4.**
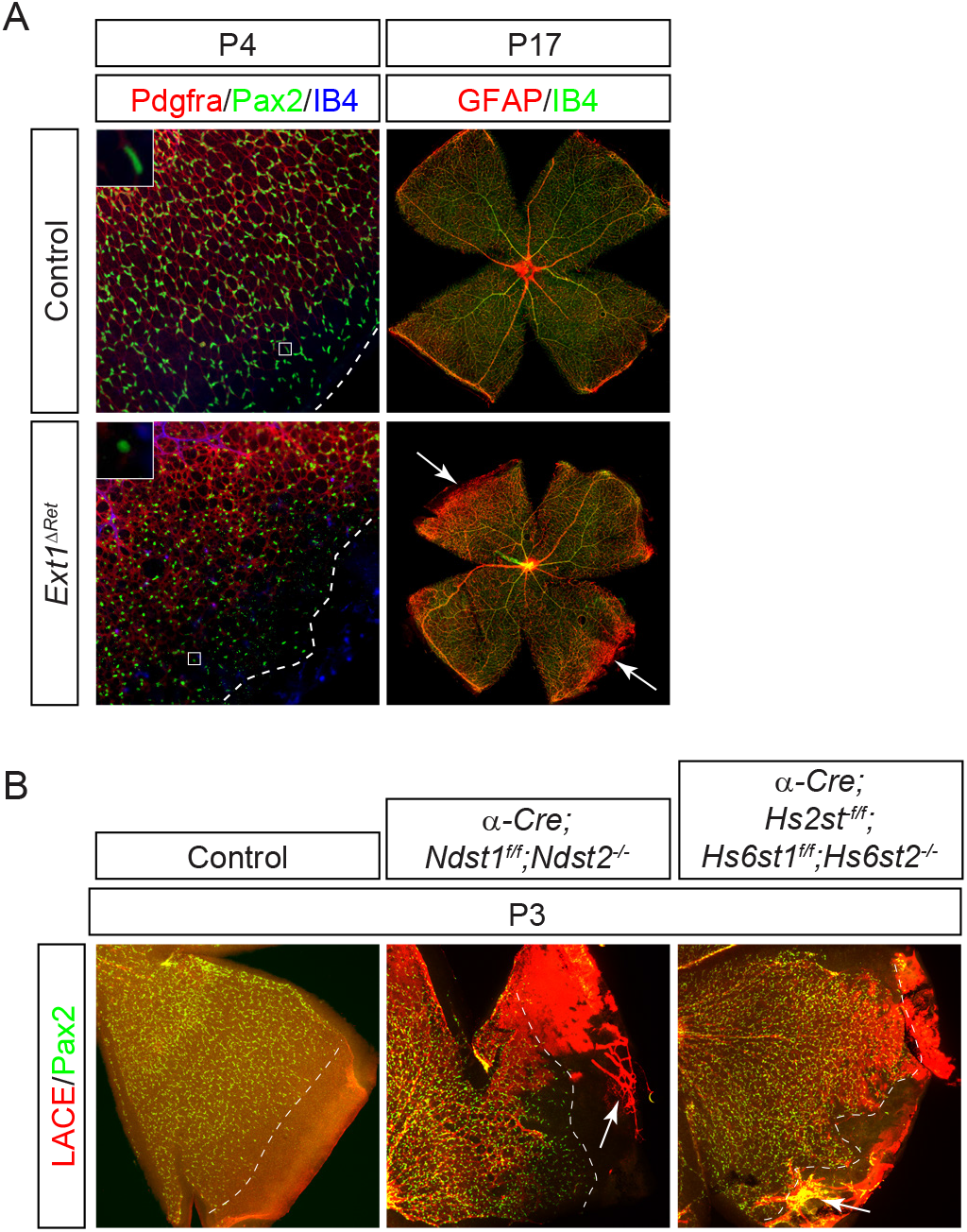
Astrocyte migration was disrupted at the peripheral retina by deletion HS biosynthesis genes. **(A)** Migratory astrocytes in *Ext1*^*ΔRet*^ mutants were uneven and sparser compared to those in controls as they approached distal retinae (dotted lines). Their nuclei shapes changed from spindle-like to round and they showed signs of stress as indicated by hypertrophic cell bodies and elevated GFAP expression (arrows). **(B)** Uneven and sparser astrocytes were also observed in HS N- and O-sulfation mutant retinae, which also displayed abnormal penetration by hyaloid vessels (arrows). Dotted lines indicate migratory fronts of astrocytes.

The HS polymer generated by Ext1 enzyme can be further modified by a number of HS sulfotransferases at N-, 2-O, 3-O or 6-O positions of glucosamine residues. To investigate the role of HS sulfation in development and distribution of retinal astrocytes, we first genetically ablated the HS N-deacetylase and N-sulfation genes *Ndst1* and *Ndst2* in the retina. As shown by the LACE assay, sulfated HS was indeed depleted in *α-Cre;Ndst1*^*flox/flox*^;*Ndst2*^*−/−*^ mutants, leading to uneven migration of astrocytes in peripheral retinae like that of *Ext1*^*ΔRet*^ mutants (Fig. 4B, dotted lines). Hyaloid vessels constitute the embryonic vasculature in the vitreous that normally regresses after birth (Lang, 1997). In *α-Cre;Ndst1*^*flox/flox*^;*Ndst2*^*−/−*^ mutants, however, there were persistent hyaloid vessels marked by strong LACE staining that became attached to HS-deficient retinae (Fig. 4B, arrows). We next deleted the HS uronyl 2-O sulfotransferase gene *Hs2st* and HS glucosaminyl 6-O sulfation genes *Hs6st1* and *Hs6st2*. Similar to *α-Cre;Ndst1*^*flox/flox*^;*Ndst2*^*−/−*^ mutants, we observed uneven migration of astrocytes and persistent hyaloid vessels in peripheral retinae in *α-Cre;Hs2st*^*flox/flox*^;*Hs6st1*^*flox/flox*^;*Hs6st2*^*−/−*^ mutants. These subtle but consistent defects suggest that sulfated HS in the retina regulates astrocyte development and migration, as well as hyaloid vessel clearance.

### Retinal HS potentiate astrocyte migration by promoting the assembly of the ILM

The ILM is essential for astrocyte migration and preventing abnormal penetration of hyaloid vessels into the retina (Edwards et al., 2010; Gnanaguru et al., 2013). Although the immunostaining of retinal sections did not reveal obvious defects in the ILM (Fig. 1B), we decided to re-examine ILM integrity more thoroughly by whole mount staining. In the majority of *Ext1*^*ΔRet*^ mutant retinal regions, the ILM appeared as a smooth layer of Laminin overlying the IB4-positive blood vessels. However, in the far periphery of mutant retina, however, we observed scattered holes in the ILM (Fig. 5A, arrowheads). Moreover, these holes were often penetrated by residual hyaloid vessels (Fig. 5A, arrow), similar to what we observed the periphery of HS N-sulfation and O-sulfation mutant retinae.

**Figure 5.**
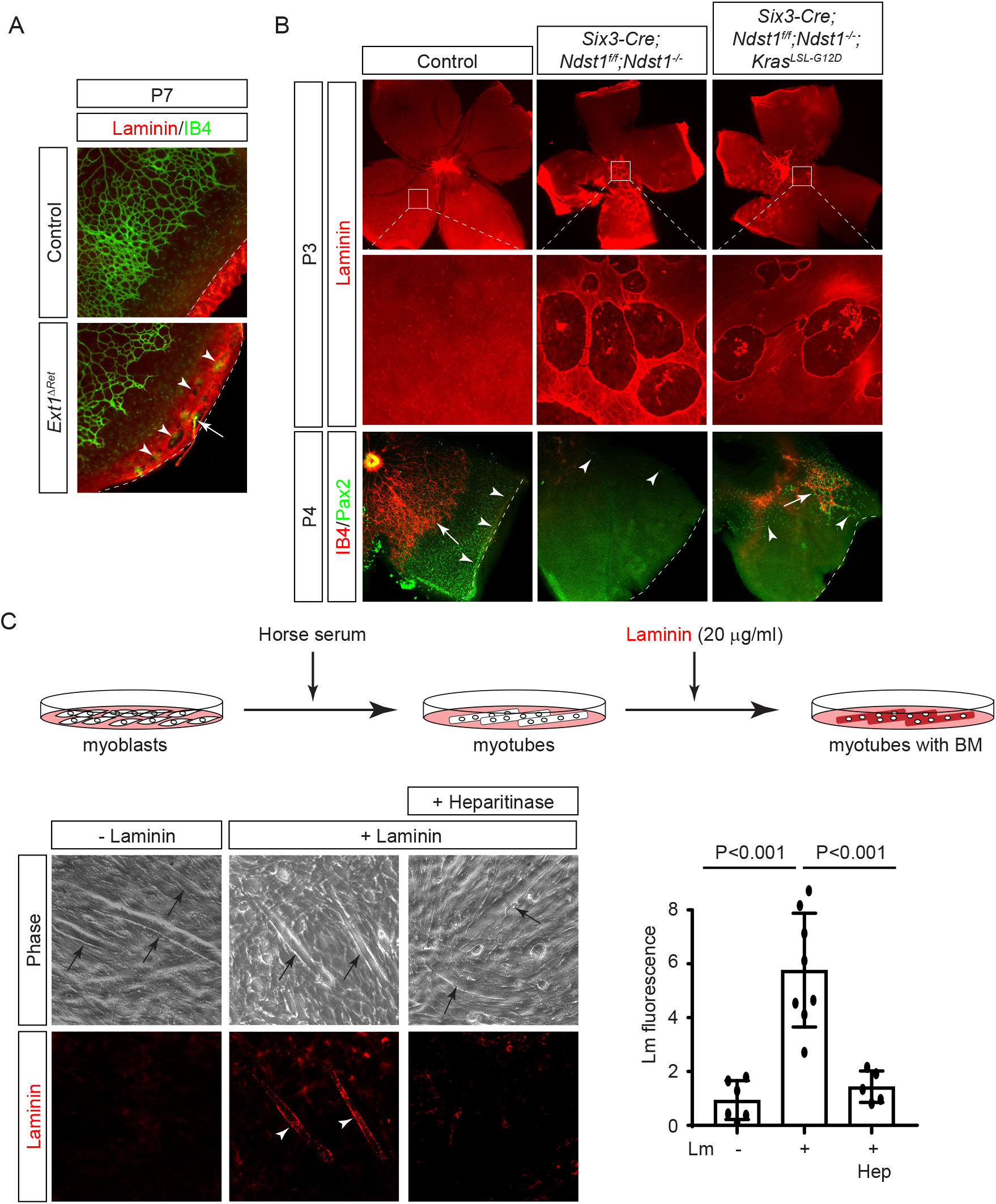
Retinal HS is required for the assembly of the ILM. **(A)** Laminin staining revealed that the peripheral end of *Ext1*^*ΔRet*^ mutant retina contained holes in the ILM (arrowheads), which were penetrated by hyaloid vessels positive for both Laminin and IB4 (arrow). **(B)** *Six3-Cre* mediated inactivation *Ndst* genes resulted in extensive ILM breaks in the central retina, which can not be rescued by constitutive Kras signaling. As a result, these mutants displayed severe defects in astrocyte migration (arrowheads) and angiogenesis (arrow). **(C)** After replacing fetal bovine serum (FBS) with horse serum, C2C12 myoblast cells fused into myotubes, which can induce exogenous Laminin to polymerize on their surface. This initiation of basement membrane assembly was blocked by Heparintinase to remove the cell surface HS. The intensities of Laminin staining on myotubes were measured using ImageJ (One-way ANOVA, *n*>4 for each group, P<0.001)

Our data revealed that the ILM defects in *Ext1*^*ΔRet*^ mutants was restricted to a narrow band at the end of the retina, whereas a wider region of HS-depleted retina showed an intact ILM. To reconcile the discrepancy between the restricted ILM defect and the extensive HS depletion, we considered the fact that retinal development follows a central-to-peripheral pattern. We hypothesized that because of the relatively late onset of *α-Cre* activity and persistent activity of previously expressed enzymes, much of HS depletion may have occurred after the formation of the ILM in the mutant retina. If this model is correct, an early Cre driver may induce much more severe disruption of the ILM. To test this hypothesis, we replaced *α-Cre* with *Six3-Cre*, which acts one day earlier in the central retina at E9.5 (Cai et al., 2013; Furuta et al., 2000). In *Six3-Cre;Ndst1*^*flox/flox*^;*Ndst2*^*−/−*^ mutants, there were indeed much larger holes in the ILM in the retina (Fig. 5B, boxes). As a result, astrocyte migration was mostly aborted, leaving only a few escaped astrocytes scattered in the retina (Fig. 5B, arrowheads). A caveat of this experiment is that HS also acts as the co-receptor for FGF signaling, which is required for development of the optic disc. However, we have previously shown that optic disc defects in both FGF receptor and *Ndst* mutants could be rescued by constitutively active Kras signaling (Cai et al., 2014). We therefore stimulated Kras signaling by inducing the oncogenic *Kras*^*LSL-G12D*^ allele and again observed extensive ILM breaches in *Six3-Cre;Ndst1*^*flox/flox*^;*Ndst2*^*−/−*^;*Kras*^*LSL-G12D*^ mutants (Fig. 5B). There were only a small number of astrocytes and associated endothelial cells in the retina, consistent with the essential roles of the ILM and HS in astrocyte migration and angiogenesis.

Study of these HS biosynthesis mutants thus far revealed a narrow time window for HS to affect ILM development, suggesting that HS may be required for the initial assembly of the ILM but not its later maintenance. To test this model, we turned to an in vitro basement membrane assembly assay (Colognato et al., 1999). In this experiment, C2C12 myoblast cells were prompted to form myotubes after replacing fetal bovine serum (FBS) with horse serum in the media (Fig. 5C, arrows). Although myotubes themselves expressed little endogenous Laminin, they were competent to assemble de novo basement membrane using exogenous provided Laminin (fig. 5C, arrowheads). However, if these myotubes were first stripped off their cell surface HS by heparin lyase treatment, Laminin polymerization was no longer detected. This result shows that the cell surface HS plays an important role in the assembly of the basement membrane.

### CS cooperates with HS to control ILM permeability

In comparison to HS biosynthesis mutants described above, our previously studied *α-Cre;Ugdh*^*flox/flox*^ mutants displayed more extensive ILM defects (Tao and Zhang, 2016). One possible reason is *Ugdh* knockout abolished biosynthesis of both HS and CS, which may have a synergistic effect on the integrity of the ILM. To investigate this possibility, we ablated the CS 4-O-sulfation enzyme C4st1 (also known as Chst11). *α-Cre;C4st1*^*flox/flox*^ (*C4st1*^*ΔRet*^) mutants did not show any obvious abnormalities in astrocyte migration and angiogenesis (Fig. 6A). As shown by Laminin staining, even combined deletion of *Ext1* and *C4st1* in *α-Cre; Ext1*^*flox/flox*^;*C4st1*^*flox/flox*^ (*Ext1;C4st1*^*ΔRet*^) mutants failed to generate more severe ILM defect than that of *Ext1*^*ΔRet*^ mutants (Fig. 6A, arrows).

**Figure 6.**
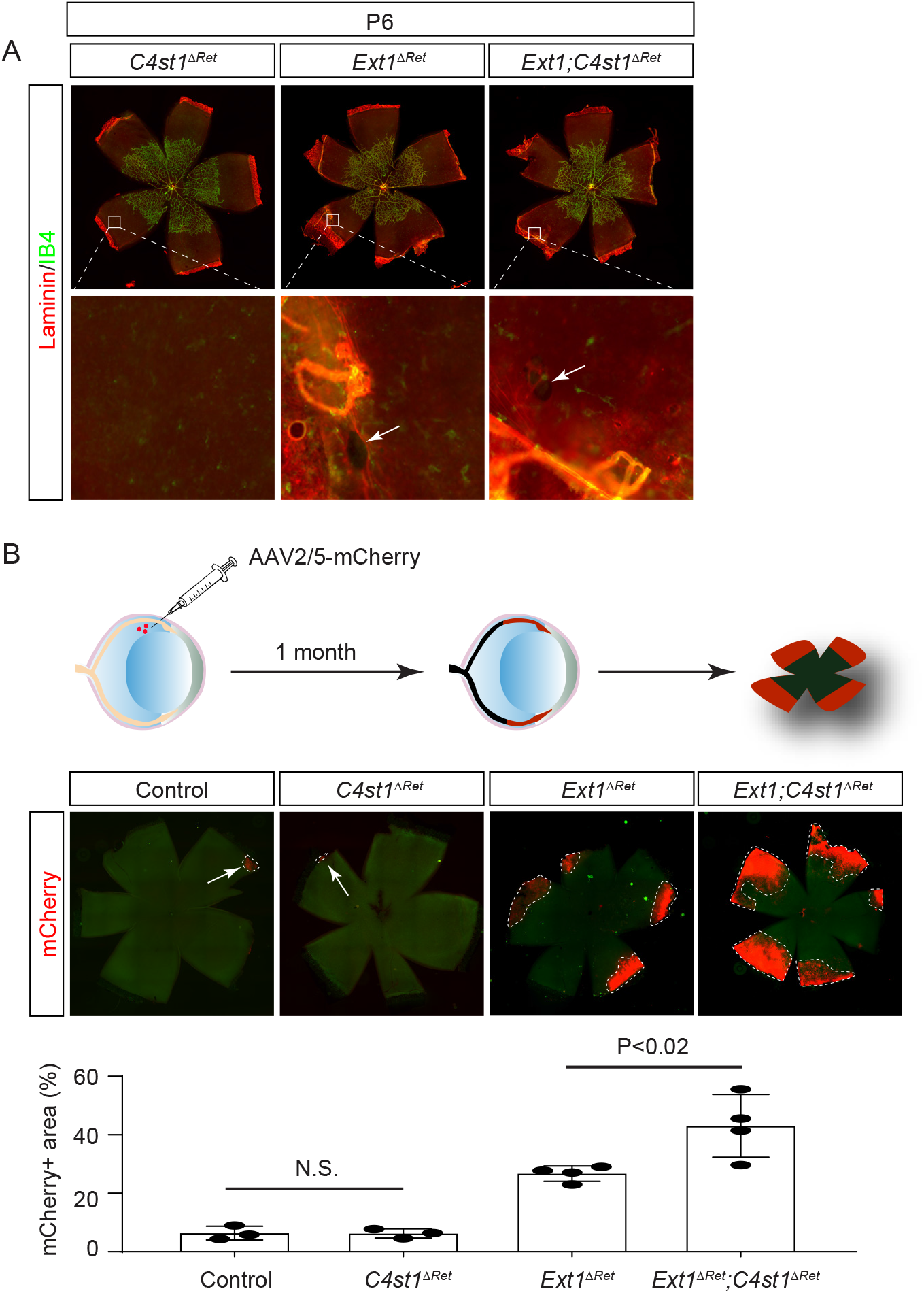
The retinal CS regulates the ILM permeability with HS. **(A)** There were no obvious ILM holes in *C4st1*^*ΔRet*^ mutant retinae and extents of ILM defects were comparable between *Ext1*^*ΔRet*^ and *Ext1;C4st1*^*ΔRet*^ mutants. **(B)**One month after intravitreal injection of AAV2/5-mCherry, control and *C4st1*^*ΔRet*^ mutant retinae expressed mCherry only in injection sites, demonstrating the integrity of their ILMs. In contrast, mCherry expression was detected in the peripheral *Ext1*^*ΔRet*^ mutant retina and significantly expanded in *Ext1;C4st1*^*ΔRet*^ mutants. The area of mCherry expression were normalized as the percentage of the entire retina. (One-way ANOVA, *n*>3 for each genotype, P<0.02, N.S., not significant)

Since deletion of C4st1 is not expected to disrupt CS function entirely, we considered the possibility that its effect on the ILM might be relatively subtle. A sensitive test for the integrity of the ILM is intravitreal injection of adeno-associated virus AAV5, which can robustly infect the retina if there are any breaches in the ILM (Dalkara et al., 2009). To detect the viral infection, we used a pseudotyped AAV2/5 virus that manifests the tropism of AAV5 but carried an AAV2 genome encoding mCherry reporter. After injection into the vitreous, this virus induced mCherry only at injection sites in both control and *C4st1*^*ΔRet*^ animals (Fig. 6B, arrows), demonstrating that the barrier function of their ILMs was intact. In contrast, *Ext1*^*ΔRet*^ mutants displayed mCherry expression in peripheral retinae (Fig. 6B, dotted lines), where we previously detected obvious holes in the ILM by Laminin staining. Importantly, mCherry-expressing domains were significantly enlarged in *Ext1;C4st1*^*ΔRet*^ mutants compared to those in *Ext1*^*ΔRet*^ mutants, matching the *α-Cre* deletion pattern in the retina. These results showed that *Ext1* and *C4st1* both contribute to the barrier function of the ILM.

## Discussion

In this study, we have explored distinctive functions of HS and CS in the neural retina. By deleting the HS polymerase Ext1 in retinal progenitor cells, we demonstrated that retinal HS was dispensable for VEGF-dependent migration of endothelial cells, but it regulates the subsequent pruning and maturation of the retinal vasculature. We also showed that retinal HS is required in a narrow time window for the assembly of the ILM, which is necessary for migration of astrocytes into the retina. Lastly, we presented evidence that deficient sulfation of the retinal CS enhanced the permeability of the ILM caused by HS depletion, revealing a novel role of CS in maintaining the barrier function of the basement membrane. These results shed light on the multifaceted function of GAGs in growth factor signaling and extracellular matrix integrity in the retina.

It is notable that our genetic ablation of *Ext1* failed to perturb the progression of the vascular front during postnatal retinal development, yet led to a more severe oxygen induced retinopathy (OIR) phenotype with more extensive capillary loss in response to hyperoxia. At first glance, this result may seem to contradict the widely accepted model that Vegfa requires interaction with the cell surface HS to maintain a chemo attractive gradient for angiogenesis (Ruhrberg et al., 2002; Stalmans et al., 2002). However, it is important to consider that our conditional knockout of *Ext1* was restricted to the neural retina, leaving the lens and the optic stalk intact as potential sources of exogenous HS. Halfter and colleagues have previously performed the elegant mouse/chick transplant experiment to show that the lens synthesizes many critical components of the ILM, including HSPGs, Perlecan and Collagen XVIII, while only Agrin is expressed solely by the neural retina (Halfter et al., 2000). This explains why weakened HS staining persisted in the ILM above *Ext1* mutant retina. In addition, the optic stalk is the source of retinal astrocytes, which migrate ahead of endothelial cells onto the retina and express their own proteoglycans. In fact, it was previously reported that an astrocyte specific knockout of *Ext1* indeed slowed down the radial expansion of the endothelial network by dampening VEGF signaling (Stenzel et al., 2011). This is in contrast to our retinal specific *Ext1* knockout, which instead disrupted the VEGF-dependent pruning and maturation of the vasculature. Therefore, HS expressed by different ocular tissues cooperate to regulate distinctive aspects of VEGF function, orchestrating the fine-tuning of vascular development. Our findings may have clinical significance as well in terms of the stability of the nascent retinal vasculature. They suggest that defects in HS expression in the retina may lead to increased vulnerability to retinopathy in premature infants (e.g. retinopathy of prematurity) without showing other obvious abnormalities in vascular development.

Another intriguing finding in our study is that α-Cre-induced abrogation of HS polymerase gene *Ext1* disrupted ILM formation, but the breach was restricted to the far periphery of the retina. This led to abnormal migration of astrocytes in this region and penetration of hyaloid vessels into the retina, phenotypes also observed in HS N- and 6-O sulfation mutants. The limited scope of ILM defect was not due to the lack of gene inactivation, as LACE staining on sections revealed a much larger region of HS depletion in *Ext1*^*ΔRet*^ mutant retinae. Instead, we note that retinal development occurs in a center-to-periphery pattern. As a result, when Halfter and colleagues performed a pulse-chase experiment to study ILM regeneration in vivo, they found the newly synthesized ILM only present in the distal retina (Halfter et al., 2000). It is conceivable that, due to the relatively late onset of *α-Cre* driver and the persistence of residual Ext1 protein, the eventual arrest of HS biosynthesis in our *Ext1*^*ΔRet*^ mutants coincided with the time of ILM assembly only at the far end of the retina. In this scenario, even if HS were lost in the more central retina already covered by ILM, only the nascent ILM above the distal retina would be affected. It suggests that the HS derived from the neural retina is only required for the initial assembly of the ILM, not its later maintenance. This model further predicts that the ILM could also become vulnerable to HS depletion in the central retina, as long as the onset of genetic knockout was early enough. Indeed, when we switched to an earlier retinal driver *Six3-Cre* to delete *Ndst* genes, we observed much larger and more numerous ILM holes in the central retina. Moreover, we showed in an *in vitro* assay that degradation of HS prevented myotubes from promoting Laminin polymerization, suggesting a general role of the cell surface HS in the basement membrane assembly. This is in contrast with the secreted HS proteoglycans Perlecan and Collagen XVIII, which are constitutional components of the basement membrane necessary for its stability (Yurchenco and Patton, 2009). These results highlight that HS plays tissue specific roles in the assembly and maintenance of basement membrane.

While we have thus far attributed the phenotype of *Ext1* knockout to the loss of HS, we also need to consider the potential gain-of-function effect of CS. It has been shown that CS biosynthesis may be upregulated in the absence of HS (Bachvarova et al., 2020). In addition, we have previously reported that deletion of GAG biosynthesis gene *Ugdh* resulted in more severe ILM defects than *Ext1* knockout, which could be due to either difference in the enzymatic kinetics between Ugdh and Ext1 or compensation by CS. To test the latter possibility, the ideal approach would be genetic ablation of CS biosynthesis. However, unlike HS whose chain elongation can be blocked by the removal of a single enzyme Ext1, there are four CS co-polymerase genes in the mouse genome that act redundantly, making it daunting to abrogate CS biosynthesis genetically (Bishop et al., 2007; Mikami and Kitagawa, 2013; Mizumoto et al., 2013). Thus, we have opted to remove C4st1, which catalyzes the important 4-O sulfation of CS, with the understanding that the function of CS will be at most partially impaired because of the expression of other 4-O-sulfotransferases. However, using the sensitive virus infection assay, we found that loss of *C4st1* enhanced the permeability of the ILM caused by *Ext1* deficiency, revealing a hitherto unappreciated role of CS in establishing the neuroretinal barrier. Breach of the ILM is the hallmark of retinopathy of prematurity, diabetic retinopathy and proliferative vitreoretinopathy. With deeper understanding of its biological function and innovative strategies to repair or stabilize its structure, the ILM may present a promising therapeutic target for preventing or curbing the harmful effect of these diseases.

## Methods and Materials

### Mice

*Ndst1*^*flox*^ and *Hs2st*^*flox*^ mice have been previously reported (Grobe et al., 2005; Stanford et al., 2010). *Chst11*^f/f^ mice were generated from *Chst11*^tm1a(KOMP)Wtsi^ chimeric mice (MMRRC) after crossing to *Flp* mice. *Hs6st1*^*flox*^ is a kind gift from Dr. Wellington V. Cardoso (Columbia University, New York, NY) (Izvolsky et al., 2008). *Ndst2*^*KO*^ is a kind gift from Dr. Lena Kjellén (University of Uppsala, Uppsala, Sweden) (Forsberg et al., 1999). *α-Cre* and *Six3-Cre* mice were kindly provided by Drs. Ruth Ashery-Padan (Tel Aviv University, Tel Aviv, Israel) and Yasuhide Furuta (M.D. Anderson Cancer Center, Houston, TX), respectively (Furuta et al., 2000; Marquardt et al., 2001). *Hs6st2*^*KO*^ and *LSL-Kras*^*G12D*^ mice were obtained from Mutant Mouse Regional Resource Centers (MMRRC) and the Mouse Models of Human Cancers Consortium (MMHCC) Repository at National Cancer Institute, respectively (Tuveson et al., 2004). All mice were maintained in mixed genetic background. The floxed animals which do not carry *Cre* transgene were used as controls. All animal procedures were performed according to the protocols approved by the Columbia University’s Institutional Animal Care and Use Committee.

### Hyperoxia treatment and Vegfa injection

Neonatal mice and their nursing mother are kept in the room air from birth through P9 or P24 before placed in a hyperoxia chamber and exposed to 75% oxygen for 7 days. Throughout this period, the P9 pups were housed with their nursing mother 24 hours a day and the oxygen level was monitored electronically. Neonatal pups (P9) were anesthetized by hypothermia over ice for a few minutes. The skin over the eye was cleaned with 70% ethanol using a cotton swab and the eyelid was opened by gently cutting along the scar tissue of the prospective eyelid with a sharp 30-gauge needle. The skin was pushed away from the eye using sterile forceps and 0.7 μl of Vegfa (100 ng/ μl) solution were injected slowly into the vitreous by inserting the glass needle in a 45° angle though the corneoscleral boundary. The eyelids were closed using cotton buds and the pups were placed on a heating mat at 37°C to recover before they were returned to their mother.

### AAV injection

Adult mice were anesthetized by intraperitoneal injection of Ketamine/Xylasine and breathing rate and toe pinch reflex were monitored to ensure full anesthesia. Sterile tropicamide and phenylephrine hydrochloride drops were applied to the eye, followed by Proparacaine hydrochloride eye drops for topical anesthesia. Lubricant eye gel was applied on each eye to prevent corneal ulcers. Using a sterile acupuncture needle, a small hole was created at the corneoscleral boundary and 0.5 μl of the AAV2/5-CMVmCherry (1×10^12^ vg/ml, VVC-U of Iowa-4220) solution was injected into the eye vitreous using a sharp microinjection glass needle. Proparacaine hydrochloride eye drops and lubricant eye gel were applied again on each eye and the mice were placed on a heating pad to recover.

### Immunohistochemistry

Histology and immunohistochemistry were performed on the paraffin and cryosections as previously described (Carbe et al., 2013; Carbe and Zhang, 2011). Whole retina fixed in 4% PFA or 10 μm rehydrated cryosections were blocked with 10% normal goat serum (NGS) for 1 hour at room temperature and incubated with primary antibody overnight at 4°C. After washing with PBS, samples were incubated with 2^nd^ fluorescent-conjugated antibody in 2% BSA for 1 hour at room temperature in dark. Isolectin GS-IB4 (IB4) conjugated with Alexa Fluor 488 (#I21411, Thermo Fisher Scientific) was applied to visualize the vasculature. Samples were washed and mounted with n-propyl gallate (NPG) anti-fading reagent and examined under a Leica DM5000-B fluorescent microscope. Antibodies used were: anti-Col IV (#AB756P, Millipore), anti-GFAP (#Z0334, Dako), anti-Laminin (#L9393, Sigma-Aldrich), anti-Pax2 (PRB-276P, Covance), anti-Pdgfra (#558774, BD Pharmingen) and anti-SMA (M085129-2, Agilent). Anti-Perlecan antibody was a kind gift from Dr. Peter Yurchenco (Rutgers University, Piscataway, NJ). At least three embryos of each genotype were stained for each marker.

The LACE assay was used to probe the *in situ* binding affinity of FGF–FGFR complexes to heparan sulfate on retina sections as previously described (Pan et al., 2006). Recombinant FGF10 and FGFR2b were obtained from R&D Systems, Minneapolis, MN.

### Basement membrane assembly assay

C2C12 cells were cultured in DME containing 10% fetal bovine serum (FBS). After switching to horse serum (5%), Laminin (#CB-40232, Fisher Scientific) was added at 100 μg/ml to trigger basement membrane assembly. The surface HS was removed by 1 U/ml Heparitinase I and III Blend from Flavobacterium heparinum (#H3917, Sigma Aldrich).

## Acknowledgements

The authors thank Peter Yurchenco for the Perlecan antibody, Drs. Wellington V. Cardoso, Yasuhide Furuta, Lena Kjellén, Ruth Ashery-Padan for mice. We also thank members of Zhang lab for discussion. The work was supported by grants from NIH (EY017061, EY018868, EY025933 and EY031210 to X.Z., and HL107150 and HL57345 to J.D.E).. The Columbia Ophthalmology Core Facility are supported by NIH Core grant 5P30EY019007 and unrestricted funds from Research to Prevent Blindness (RPB). C.T. is a recipient of Jonas Scholar award. X.Z. is supported by Jules and Doris Stein Research to Prevent Blindness Professorship.

